## Supplemental Material for "Continuous endosomes form functional subdomains and orchestrate rapid membrane trafficking in trypanosomes"

Membrane recycling, African trypanosomes, Endocytosis, Rab proteins, Tokuyasu, Tomography

### Supplementary Figures

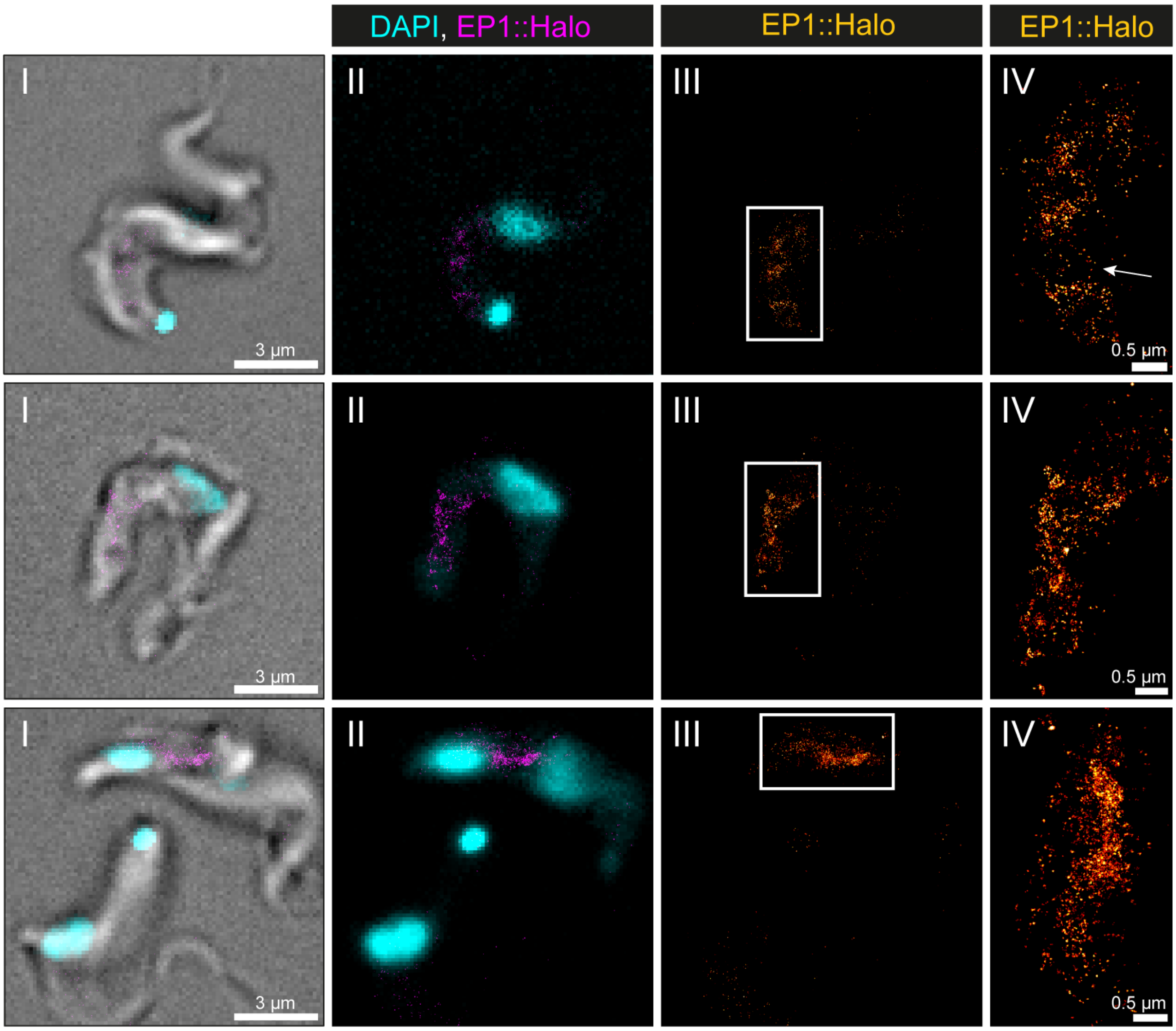


**Supplements 1. Visualisation of the endosomal system in *T. brucei* by direct stochastic optical reconstruction microscopy (*d*STORM).** A transgenic EP1::HaloTag expressing bloodstream form cell line was generated and used to visualise the endosomal system. Cells were labelled with a HaloTag ligand and fixed with 4 % formaldehyde. Shown are a merge of transmitted light, DAPI staining and the EP1::HaloTag signal (I), a merge of DAPI staining and the EP1::HaloTag signal (II), the EP1::HaloTag signal (III) and a magnified view of the EP1::HaloTag signal (IV).

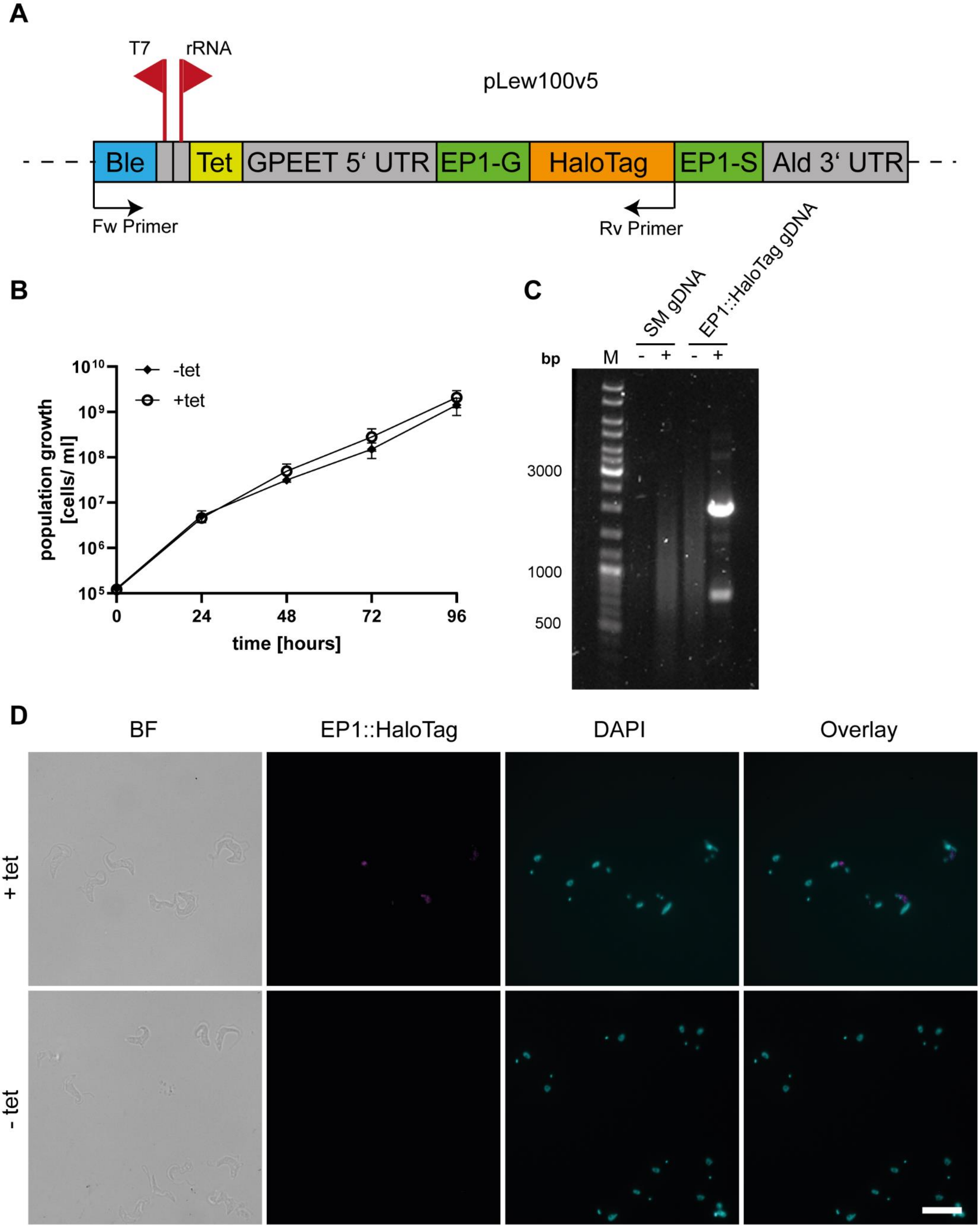

A
T7
rRNA
pLew100v5
Ble
Tet
GPEET 5‘ UTR
EP1-G
HaloTag
EP1-S
Ald 3‘ UTR
Fw Primer
Rv Primer
B
-tet
+tet
population growth [cells/ ml]
time [hours]
C
SM gDNA
EP1::HaloTag gDNA
bp
M
3000
1000
500
D
BF
EP1::HaloTag
DAPI
Overlay
+ tet
- tet


790

**Supplements 2. Validation of the generated EP1::HaloTag cell line.** (A) Schematic of the tagging construct used for the inducible, ectopic expression of the Halo-tagged version of EP1. The bleomycin (Ble) resistance is driven by the T7 promoter while the expression of the EP1::HaloTag construct is driven by a ribosomal RNA promoter that is regulated via a tetracycline repressor binding site (Tet). The EP1 construct is flanked by a GPEET 5'UTR and an Aldolase 3'UTR. Forward (Fw) and reverse (Rv) primer used for validation of integration by PCR are indicated. (B) Growth curve of the uninduced (- tet) and induced (+ tet) EP1::HaloTag cell line. Three independent clones were counted over the time of four days to confirm that the expression of the construct was not interfering with the overall cell vitality. (C) Integration PCR with parental cell line (SM) and EP1::HaloTag gDNA as template. Reactions were run with (+) and without (-) template DNA. As the plasmid would integrate into an unspecific ribosomal RNA spacer region, primers were designed to amplify the antibiotic resistance together with the EP1::HaloTag protein sequence. No product was visible in the parental cell line, while a main product of the expected size (2000 bp) was visible in the EP1::HaloTag cell line. (D) Immunofluorescence of uninduced and induced EP1::HaloTag cells. Cells were labelled with Halo ligand TMR to visualise the EP1 signal and DAPI to show the nucleus and kinetoplast. Additionally, a brightfield image (BF) and an overlay of the fluorescence channels are given. Without induction, no signal of the Halo-tag was visible. After induction, a clear Halo-tag signal was visible which also corresponds to the anti-EP1 signal. The main localisation of the signal is between the kinetoplast and the nucleus. The gene expression is heterogenous in the population as not all cells show a strong fluorescent signal. Scale bar: 10 µm.

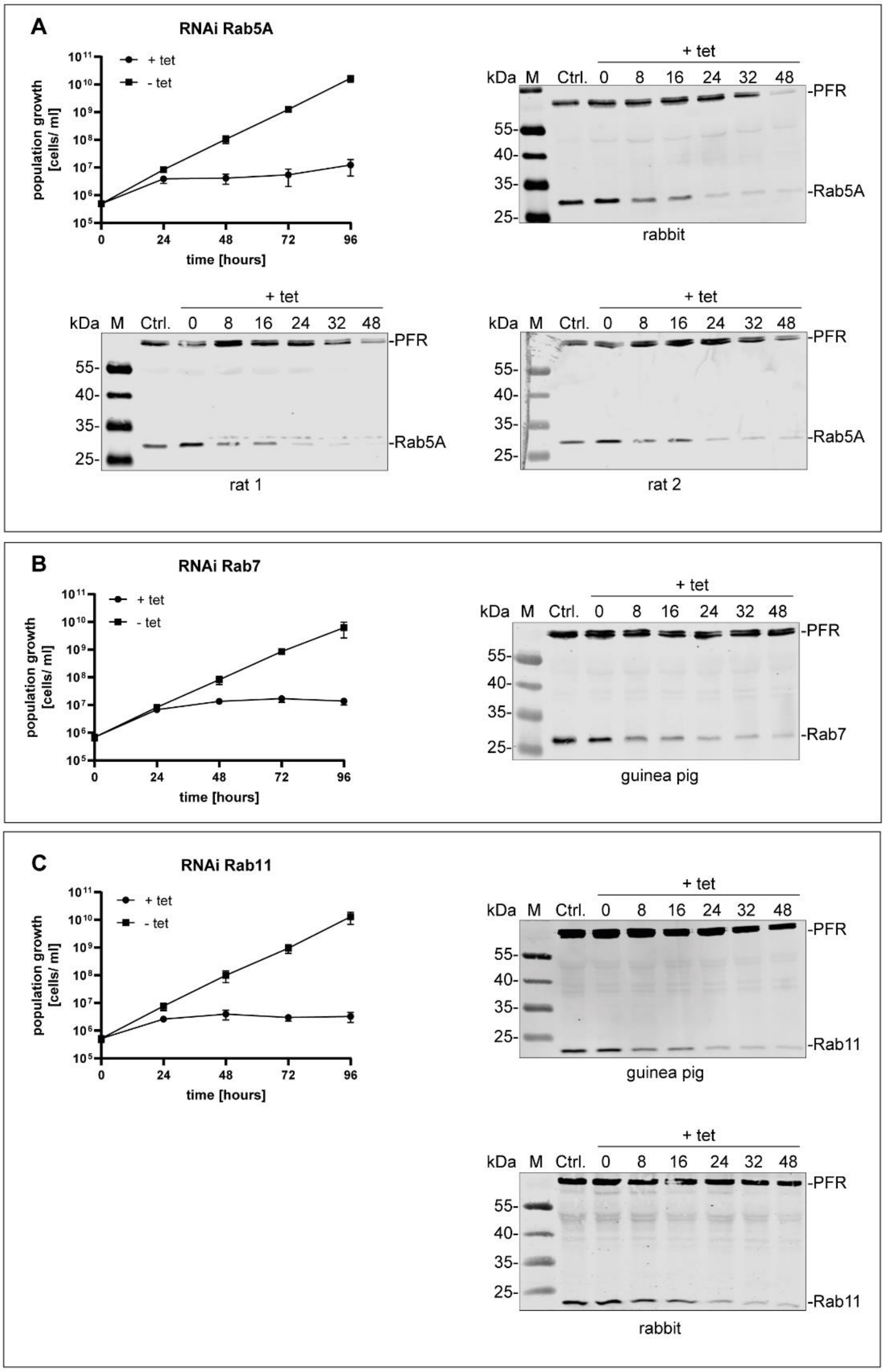


**Supplements 3. Validation of different anti-TbRab antibodies.** Antibodies against (A) TbRab5A, (B) TbRab7, (C) TbRab11 were generated in different animals and validated by immunoblotting using protein samples of RNAi cell lines. Each panel shows the cumulative growth profile of the corresponding RNAi cell line. Three independent clones were analysed for 4 days in the presence (+ tet) or absence (- tet) of tetracycline. In addition, Western blot analysis of the protein expression of the different TbRab in the corresponding RNAi cell line 0, 8, 16, 24, 32 and 48 hours after tetracycline induction. The parental 2T1 cells served as a positive control and the TbPFR1,2 (PFR) proteins served as loading control. For each TbRab protein different numbers and species of animals were immunised. Depicted are those that gave antisera with compatible antibodies.

#### Supplementary Movies

**Movie 1: Tomographic reconstructions of *T. brucei*.** The movie is available online and includes annotations for better understanding.

**Movie 2: Tomographic reconstructions of *T. brucei*.** Cells were chemically fixed before further processing.

**Movie 3: Tomographic reconstructions of *T. brucei*.** Cells were chemically fixed before further processing.

**Movie 4: Tomographic reconstructions of *T. brucei*.** Cells were chemically fixed before further processing.
